# Induction of Kanizsa contours requires awareness of the inducing context

**DOI:** 10.1101/050526

**Authors:** Theodora Banica, Dietrich Samuel Schwarzkopf

## Abstract

It remains unknown to what extent the human visual system interprets information about complex scenes without conscious analysis. Here we used visual masking techniques to assess whether illusory contours (Kanizsa shapes) are perceived when the inducing context creating this illusion does not reach awareness. In the first experiment we tested perception directly by having participants discriminate the orientation of an illusory contour. In the second experiment, we exploited the fact that the presence of an illusory contour enhances performance on a spatial localization task. Moreover, in the latter experiment we also used a different masking method to rule out the effect of stimulus duration. Our results suggest that participants do not perceive illusory contours when they are unaware of the inducing context. This is consistent with theories of a multistage, recurrent process of perceptual integration. Our findings thus challenge some reports, including those from neurophysiological experiments in anaesthetized animals. Furthermore, we discuss the importance to test the presence of the phenomenal percept directly with appropriate methods.

## Introduction

What role does conscious processing of the environment fulfill and how much processing occurs in the absence of awareness? It is self-evident that much of the internal bodily functions and the learned motor behaviors, such as walking or driving, operate mostly without awareness. But for processing through the classical senses, like vision, there have been widely discrepant findings on how much stimulus processing can occur and how it affects decision-making when the subject is unaware of the stimulus. Moreover, the approach to be used when studying unconscious stimulus processing has also been subject of controversy [1].

Several experiments suggest that the effect of contextual stimuli within a target, such as the percept of visual illusions or adaptation effects, persists even when participants are unaware of the presented contextual information [2–6]. The use of continuous flash suppression (CFS), in which a dynamic, high-contrast stimulus is presented to one eye to suppress the stimulus in the other eye from awareness, has become a popular way to probe unconscious stimulus processing [7]. Using this procedure it has been claimed that the perception of physical facial attributes [8], the complex analysis of naturalistic scenes [9], and even linguistic processing and arithmetic can be performed without awareness [10]. However, several of these findings have recently failed to be replicated and were challenged on theoretical grounds [11–13].

Neuroimaging experiments showed that while both simple and more complex stimuli have a neural signature in the visual cortex under masking conditions [14–16], the encoding of unconscious stimuli appears to be qualitatively different. Not only is the overall response to unconscious visual stimuli weaker [15] but coupling between different stages in the visual processing hierarchy is also reduced [17] and the information content differs [16,18,19]. In particular, the response to these stimuli is more variable [20], and also localized in more posterior regions than to conscious stimuli [18]. . One reason for this variability could be that only simple positional or geometric information is processed in the absence of awareness, but that more complex abstraction and perceptual integration requires consciousness Under this hypothesis, the neural encoding of stimuli is noisy because local stimulus interactions are preserved but abstract and therefore coherent representations are disrupted. Specifically, we used shape stimuli that were either defined by the position or the orientation of simple image elements. We demonstrated that when such stimuli were rendered invisible using fast counter-phase flicker (at 120Hz) they could speed up performance on a shape discrimination task of conscious stimuli [21]. Critically, this priming effect was only present for stimuli sharing the *same* positions, or — if oriented elements were used for priming — the positions along the path implied by the elements. We observed no priming by invisible primes if the test stimulus was smaller than the prime stimulus. This suggests that invisible priming operated locally, possibly in retinotopic space in early visual cortex, but no abstract integration of individual elements into the concept of a shape occurred without awareness.

We further tested whether two brightness illusions manifest when the inducing context is rendered invisible by means of CFS [22]. We found that masking the context (a smooth gradient in luminance) had little impact on simultaneous brightness contrast of two targets that were unmasked. In a stark dissociation, participants could not discriminate the orientation of an illusory contour, defined by a Kanizsa triangle, when the inducing context (the ‘Pacman’ shapes whose open segments define the corners of the triangle) were masked selectively by CFS. This could indicate that the generation of the illusory contour percept occurs at a later stage of visual processing than simultaneous brightness contrast, either in terms of the visual hierarchy or in the latency of processing. These findings were also consistent with previous reports that illusory contours are not perceived when the inducers are suppressed from awareness during binocular rivalry [23].

These findings challenge some previous reports, using psychophysical tests in healthy volunteers [24] and even neurophysiological experiments in anesthetized animals [25] that suggested that illusory contours could be formed in the absence of awareness. However, none of these previous studies specifically tested whether the experimental participants actually *perceived* an illusory contour. Similarly, neuropsychological studies in neglect patients [26–29] suggested that illusory contour processing occurs when part of the inducing context is placed in the blind hemifield. However, this also does not conclusively support the assertion that contours are formed in the absence of any contextual awareness.

However, two issues burden the interpretation of previous experiments that show no evidence of illusory contours when the inducers are masked. First, there is evidence that illusory contours are processed by binocular neurons in early visual cortex [30–33]. It does in fact seem unsurprising that the mechanisms inducing illusory contours at least partially overlap with those for segmenting surfaces in depth [34]. When retinal disparity implies that inducers are at different depths from the background, the visual system not only produces the percept of illusory contours but the surface bounded by illusory contours is also perceived in stereoscopic depth [31]. Thus when binocular processing is disrupted or overwhelmed by a dichoptic mask or binocular rivalry, the illusory contour percept is also broken. Another recent study further complicated this situation by finding that Kanizsa shapes broke through CFS faster than control stimuli [35]. Leaving aside conceptual issues with the time-to-emergence paradigm, it is imperative that the dependence of illusory contours on awareness must be confirmed using masking methods other than CFS.

A second confound with these previous studies [22,23] is that even if participants perceived an illusory contour, this percept was far less salient than real luminance contours, as it may have been obscured by the dominant masking stimulus. The addition of a simple control condition could remedy this problem: One experimental stimulus should be a *real contour* defined by a subtle luminance contrast that mimics the illusory contour percept as closely as possible. If participants can detect and discriminate this stimulus but are unable to do so for the illusory contour condition, this indicates that the *illusory percept* is indeed disrupted specifically, rather than the more general ability to detect subtle stimuli.

In the present paper, we carried out two experiments to address these confounds and answer the question of whether Kanizsa contours are formed when inducers are not consciously perceived. In experiment 1, we used a similar design as in our previous study [22]. Here, participants were asked to discriminate the orientation of a Kanizsa triangle. However, instead of CFS we employed a temporal masking method to render the inducers invisible. Moreover, we included a control condition in which a real, luminance-defined contour was present. Because this masking method relies on very brief stimulus durations, in experiment 2 we presented long (500ms) stimuli rendered invisible by means of fast counter-phase flicker [3,21]. This is critical because the formation of illusory contours arise comparably slowly [36–38] and thus may be disrupted by a fast temporal masking technique. In addition to this, previous research demonstrated that the presence of illusory contours boosts participants' ability to discriminate the position of a tiny target [23,39], providing a specific test of whether the participant in fact perceives an illusory contour or not. We therefore measured the ability of a group of participants, who were well trained at psychophysical tasks, to discriminate the position of a dot target for Kanizsa and control stimuli presented with or without masking.

## Materials and Methods

Both experiments were carried out at the UCL Department of Experimental Psychology. Procedures adhered to the Declaration of Helsinki. Ethical approval for this study was obtained from the UCL Research Ethics Committee and all participants gave written, informed consent.

Participants in experiments 1 were recruited among the UCL student population. In experiment 2 we recruited participants who were familiar with psychophysical tasks. All Participants had normal or corrected-to-normal visual acuity.

All experiments were conducted in a dark, sound-attenuated room. For the duration of the experimental sessions, participants were asked to stabilize their head on a chin rest located at a fixed distance of 48cm from the stimulus presentation screen where stimuli were presented to them binocularly.

Stimuli were generated by a computer and presented on a 22-inch Samsung SM2233RZ LCD monitor at a resolution of 1680 x 1050 pixels. Screen refresh rate in experiment 1 was set to 60Hz. In experiment 2 it was set to 120Hz. The experiment was controlled and behavioral responses were recorded using MATLAB (The Mathworks, Inc.) and Psychtoolbox 3 [40] using a standard keyboard.

### Experiment 1

The experiment comprised two tasks: the first, henceforth called ‘Kanizsa’ task, investigated whether participants can perceive illusory contours without awareness of their inducers. The second task, the ‘Visibility’ task, assessed the effectiveness of the masking technique directly.

#### Study design

Like our earlier experiment using continuous flash suppression, this experiment aimed to measure the perception of illusory contours in a direct manner. We implemented a 2×3 design with visibility (invisible, visible) and type of stimulus (real, illusory, control) as within subject factors.

Every participant completed two different tasks. Each task comprised 25 blocks of trials, where one block consisted of 24 trials in the Kanizsa task and 8 trials in the visibility task. Across one task, each condition appeared 100 times. The visibility task only comprised the visible and invisible control conditions.

Participants made behavioral responses in a forced-choice design by button press (left or right arrow) on a standard computer keyboard. Each trial required either a left or a right response and each response type appeared twice per block for each condition. Conditions were selected pseudo-randomly for every trial but were counterbalanced over each block.

#### Participants

Seventeen (13 female; age range: 18-29, mean age: 23.8±2.5) normal, healthy participants took part in the experiment. An additional two participants were tested but they failed to discriminate the real luminance contour under masking conditions and were therefore excluded from further analysis (see more details below). All participants were unaware of the experimental hypothesis.

#### Stimuli

Stimuli were created by placing four discs (inducer elements, diameter=2.2°) in the configuration of a square (width=4.3°). The configuration of these discs was centered on fixation. As both standard edge type and a number of line-end inducing elements have comparable efficacy in the clarity with which illusory contours are perceived by participants [41,42], here the discs were defined by partial concentric circles. Each of the four lines forming the circles had a width of approximately 0.07° with a luminance of 0.6 cd/m^2^. The positioning of the gap within the circles gave rise to the percept of a Kanizsa triangle (Figure 1A). Thus, a number of line-end inducing elements gave rise to perception of illusory contours [43].

We also included a real luminance condition. Here, the stimuli did not contain any discs but instead there was a triangle defined by a real but subtle luminance contrast at the exact location where the Kanizsa triangle would be perceived in the illusory condition (Figure 1B). We reasoned that if participants were unable to discriminate the orientation of this subtle luminance edge when the inducers were masked, this implied that they would be unable to detect any illusory contour that could have formed without awareness of the inducers. Therefore, we removed two participants whose discrimination performance for this condition was at chance levels from any further analyses.

Control stimuli did not form any triangle and were created by altering the orientations of the inducers by a systematic rotation of 180° (Figure 1C). This condition was somewhat unnecessary because the dependent variable, accuracy for discriminating the orientation of the triangle, was orthogonal to the condition and for these control stimuli there was no strictly correct answer in this discrimination task. The “correct” responses in this condition were dummy coded so that the stimulus-response mapping matched that for the equivalent illusory triangle stimuli before rotating the inducers. We included this condition as catch trials - participants should be guessing here because there was no triangle to discriminate. Thus it provided information on whether participants could indeed perceive the triangle contours or whether they adopted a strategy of matching the inducer orientation to feedback.

The background had a luminance of 76.6 cd/m^2^ and real triangles were defined by a subtly greater luminance of 86.9 cd/m^2^. The mask was only used in the invisible conditions and was designed as a square configuration of four black discs (0.6 cd/m^2^), each containing four white concentric circles (230 cd/m^2^). It was wide enough to cover all four Pacmen in the illusory and control conditions (Figure 1D). The masking technique consisted of a sequence of three frames which was repeated three consecutive times: At first the mask appeared on the screen, followed by the appearance of the stimulus and a final blank screen. In the visible conditions, a blank grey screen replaced the mask frame.

**Figure 1.**
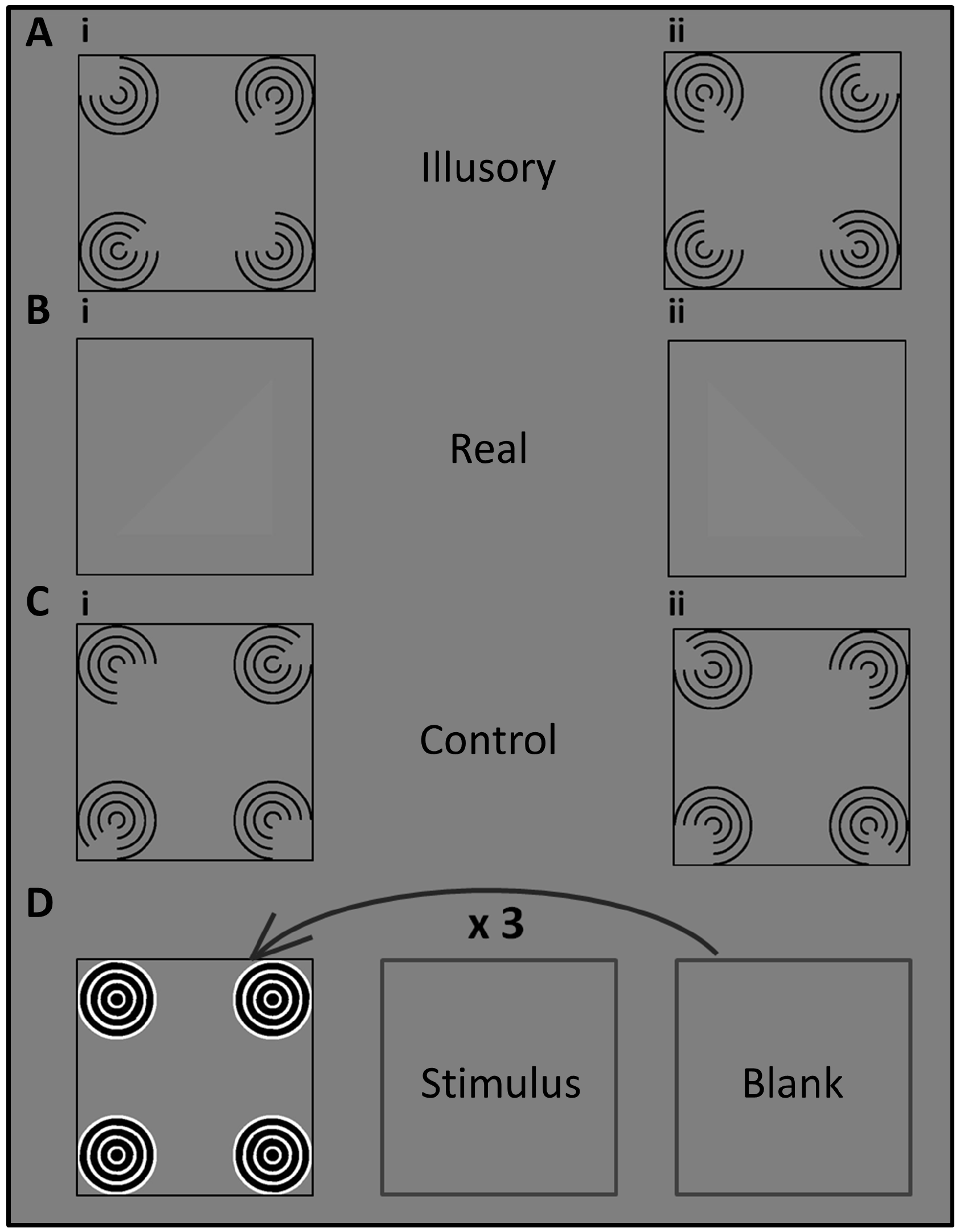
**Illustration of the stimuli**. A) (i) Kanizsa triangle pointing right (ii) Kanizsa triangle pointing left. B) (i) Real triangle pointing right (ii) Real triangle pointing left. C) Control stimuli were created by a systematic 180° rotation of the individual Pacmen. D) Illustration of the masking procedure in the invisible conditions: A mask-stimulus-blank screen frame sequence was repeated three times. In the visible conditions a blank screen replaced the mask.

#### Procedure

*Kanizsa task:* At first, participants were instructed that they would see a triangle appear on the screen. They were asked to judge whether its hypotenuse was tilted clockwise or counter-clockwise from vertical by pressing the corresponding response key. We explained this task to them as a decision whether the right angle of the triangle was pointing to the left or to the right but they were told explicitly to *judge the contours of the triangle,* in particular the long hypotenuse extending through the center of the stimulus display. Because we wished to keep the contrast between real triangle and the background as low as possible, we trained participants on the visible and invisible real triangle condition until they were able to detect them correctly.

Participants were instructed to fixate a small black dot (0.2° wide) that was present in the center of the screen throughout the experiment. On each trial, the fixation dot was displayed alone for 500ms. This was followed by a sequence of three frames that defined whether the condition was a visible or an invisible one. In the invisible condition a 300ms mask followed the fixation period. Subsequently, the stimulus appeared on the screen for one frame of approx. 16.7 milliseconds (ms), immediately followed by a blank screen that was shown for one frame as well. This mask-stimulus-blank sequence was repeated three times before a second and final post-stimulus blank screen was presented until participants gave their response. Figure 2A shows the general paradigm for the experimental procedure.

*Visibility task:* We further tested whether participants indeed did not consciously perceive any contextual information of the stimuli in the invisible condition. For this purpose, the visibility task assessed the effectiveness of the masking technique by measuring whether participants could consciously discriminate the inducer elements. Participants were asked to judge whether a right angle was presented in the left or right bottom inducer. The timing of the trial sequence in this task was identical to the Kanizsa task, with one exception: only control stimuli were used (Figure 2B).

**Figure 2.**
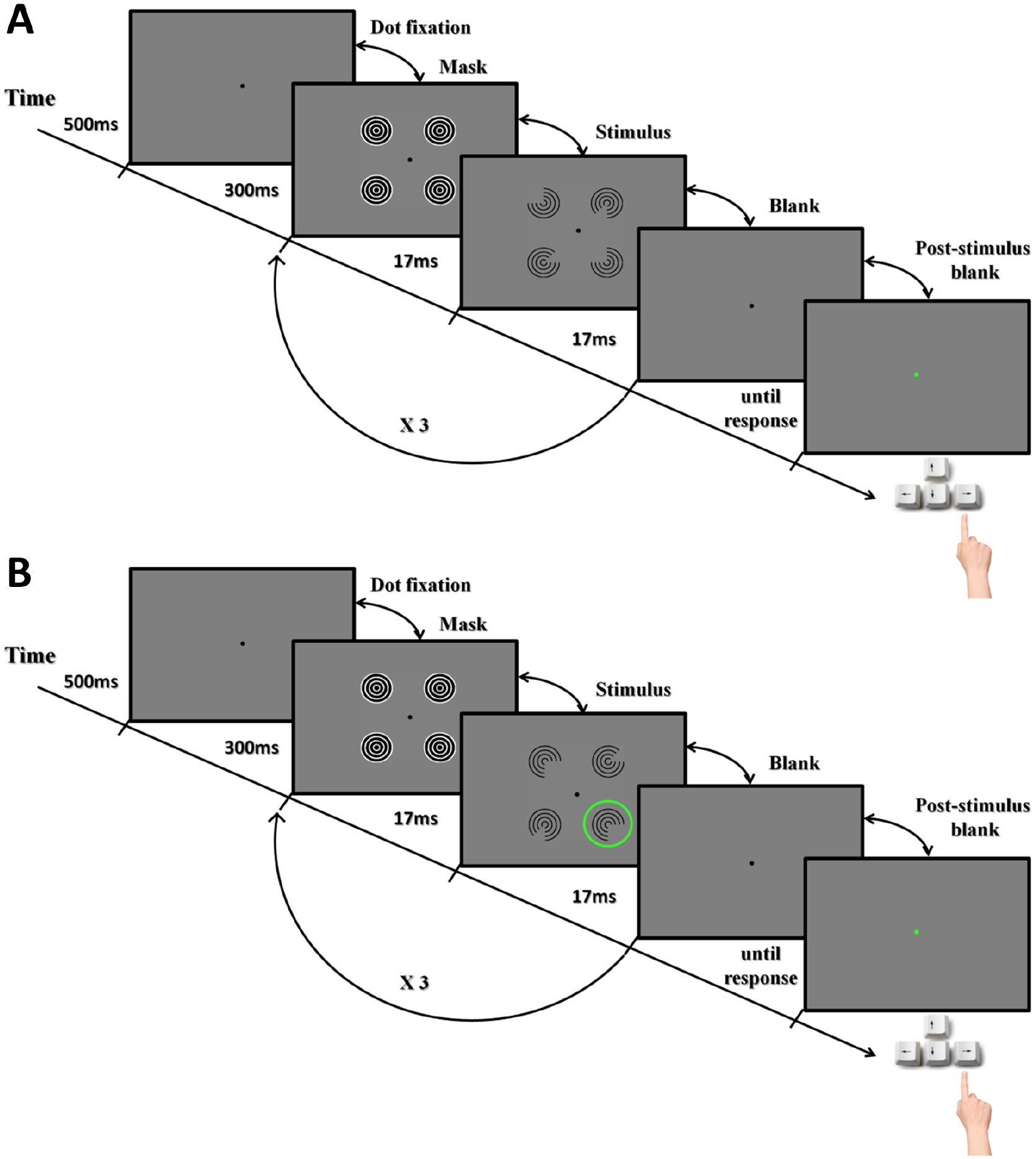
**Illustration of the trial sequence in experiment 1**. A) Kanizsa task: Each trial was composed of five frames: fixation dot, mask, stimulus, blank, and post-stimulus blank. After the participant's response, the fixation point provided feedback for 100ms (green: correct; red: incorrect). The duration of each frame is shown on the time-line. Note that in the visible conditions, a blank screen replaced the mask. B) Visibility task: Trials proceeded in the same way as in the Kanizsa task except that only control stimuli were being shown and participants judged whether the right-angle gap was in the bottom left or right inducer (indicated by the green circle, which was not present in the actual stimuli).

#### Data analysis

Performance in both tasks was defined in terms of mean proportion of accurate responses in each condition. Initially, we conducted binomial tests at the individual level to quantify how many participants performed significantly above chance. A condition for any participant's data to be considered in the group analysis of this experiment was that their performance to the invisible real stimulus condition was significantly above chance (0.5). This is because the features necessary to distinguish the contours of the real triangle were not masked and if a participant could not perform the task for this particular stimulus, any test of the perception of illusory contours would be redundant.

One-sample t-tests were carried out at the group level for each condition individually to assess whether the participants' level of performance was significantly above chance (>0.5). In addition to traditional frequentist statistical tests, we also quantified the evidence for or against the hypothesis that participants could discriminate the stimuli by calculating a Bayes Factor using a default Cauchy prior with scaling factor 0.707 for the alternative hypothesis [44]. For conditions with performance near chance levels, this enabled us to also quantify how strongly the evidence supported the null hypothesis that participants were actually guessing.

### Experiment 2

This experiment only comprised one task that tested the percept of illusory contours by measuring participants' threshold on a spatial localization task. Such a manipulation has been used successfully in previous experiments [23,39]. While it is an indirect test, the contour aids performance on the spatial localization task. This provides independent evidence about whether participants perceived any illusory contours and therefore helps to address confounds with measuring the percept directly as we did in experiment 1. Pilot experiments using a task in which we directly measured the illusory percept (the same task as experiment 1 but with the long-lasting masking technique employed in experiment 2) could encourage participants to pay close attention to the masked context and thus reduce the effectiveness of the masking procedure. This would mean that participants use residual awareness of the inducers to perform the task instead of actually making a perceptual judgment of the illusory contour.

Moreover, because the masking procedure in experiment 1 used very brief presentations of the invisible Kanizsa stimuli (16.7ms) while these were much longer (300ms) for the visible ones, in experiment 2 we used a different masking procedure: stimuli were defined by sinusoidal gratings which reversed contrast polarity at 120Hz. This method can effectively render stimuli invisible for prolonged periods so that participants only perceive a grey screen. Previous research suggests that stimuli masked in this way are processed in early visual cortex [3]. Moreover, we showed that stimuli rendered invisible by this method could induce local priming effects on a shape discrimination task [21].

#### Participants

Five normal, healthy participants (3 female; age range: 24-37; mean age: 30.2) completed this experiment, including one of the authors (DSS). All participants were experienced with psychophysical experiments. While the results from a larger subject base might generalize more to a broad population, we reasoned that only precise measurements of participants' visual performance would afford interpretable data. Unlike in the first experiment, in experiment 2 we measured how position discrimination thresholds changed between experimental conditions. Naive, untrained participants from the general population would be more likely to contribute noisy data and this might obscure potential subtle perception of illusory contours in the masked condition.

#### Stimuli

We generated a Kanizsa shape by presenting four Gabor patches (sinusoidal gratings with wavelength 0.33° visual angle convolved with a Gaussian with standard deviation 0.8° at 30% contrast) in the locations of the four corners of a square with a side-length of 8.2°. We turned these patches into Pacmen by setting a right-angled region of each patch to zero contrast (uniform background grey). This stimulus thus described an illusory square (Figure 3A). In the control conditions, the Pacmen were rotated by 180° so that the corners faced outward breaking the illusory percept (Figure 3B). In the real luminance control, we did not present any Gabor patches but instead a square region with a subtle luminance contrast (54 cd/m^2^) relative to the background (Figure 3C). The orientation and phase of each Gabor patch was randomized in each trial. Finally, we created a mask stimulus by overlaying 24 Gabor patches in the four locations. These patches were presented at 5% contrast and covered the full range of orientations in equal steps of 15° but their phases were randomized. This resulted in a patchy pattern without any obvious orientation cue (Figure 3D). The mask pattern was also generated anew in every trial.

These stimuli were presented in the center of the screen. The target stimulus was a tiny dark grey dot (diameter: ~0.1°) which could appear somewhere along the horizontal meridian, either near the right or the left vertical boundary of the square.

The square stimuli were either invisible or visible. In the invisible condition, the gratings reversed contrast polarity at 120Hz. In the visible conditions and the real luminance control, every even-numbered frame was uniform grey (except for the fixation dot).

**Figure 3.**
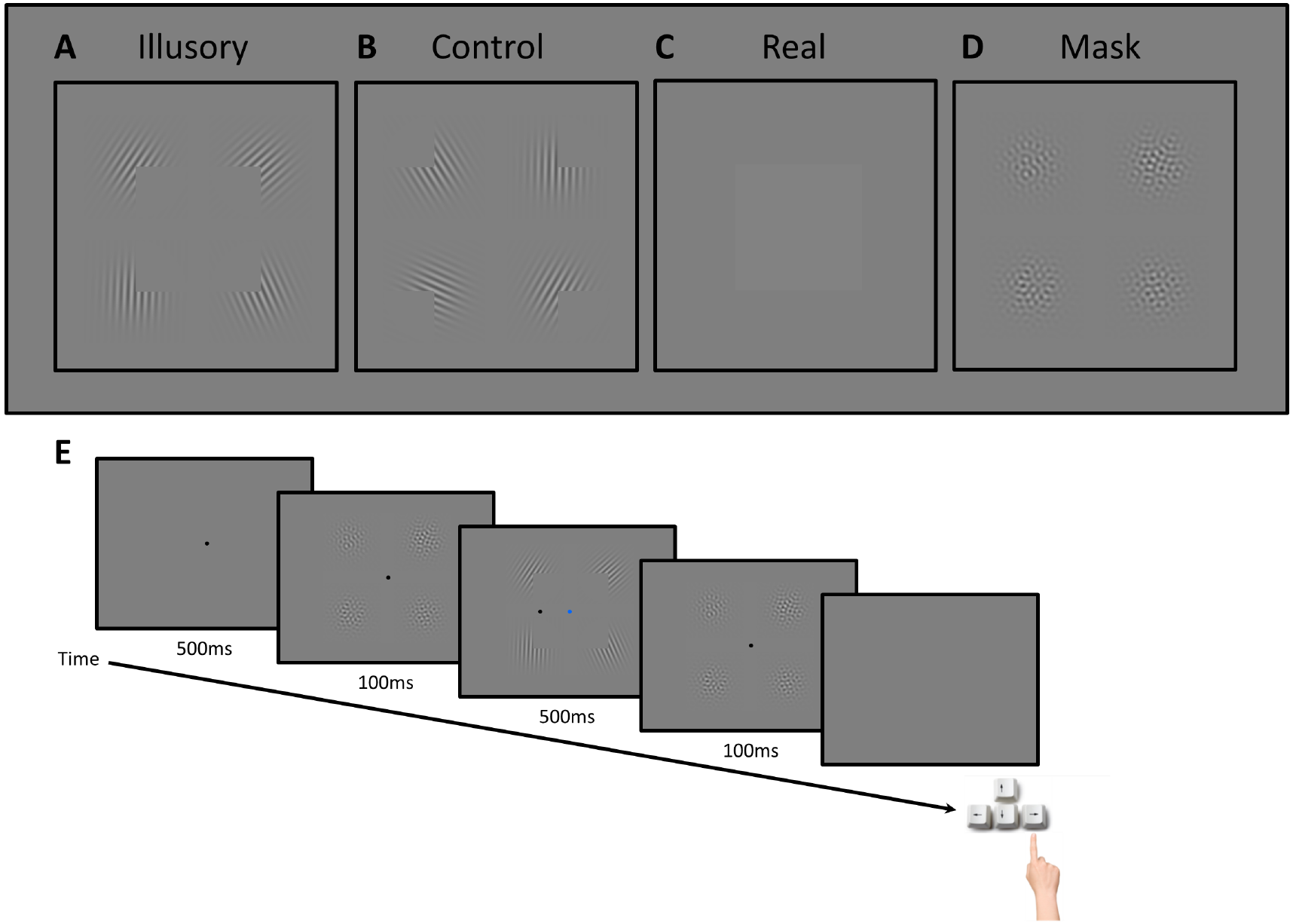
**Stimuli (A-D) and trial sequence (E) in experiment 2**. During the stimulus interval either an illusory Kanizsa square (A), a control stimulus with rotated inducers (B), or a real luminance square without any inducers (C) was presented. In visible trials every odd-numbered video frame at 120Hz contained the stimulus while every even-numbered frame contained only a blank screen with the blue fixation dot. In invisible trials, both frames contained the Gabor patches for A and B but their contrast polarity alternated between frames. For real luminance stimuli the even-numbered frames always contained a blank screen. D) A mask stimulus preceded and followed each stimulus interval. E) Each trial started with 500ms of fixation, followed by a 100ms mask, the 500ms stimulus interval, another 100ms mask, and finally a blank screen without a fixation dot that remained until participants gave their response. Their task was to locate the small target dot in the stimulus display and decide whether it appeared to the left or the right of the vertical boundary of the square (here, the correct response is right).

#### Procedure

Figure 3E depicts the sequence of an invisible Kanizsa trial in experiment 2. Each trial started with 500ms of a blank grey screen that only contained a black fixation dot (diameter 0.26°). This was followed by a 100ms presentation of the mask and then 500ms of the stimulus, after which another 100ms mask interval was presented. Then the screen turned grey and the fixation dot was removed, indicating that participants could give their behavioral response.

During the stimulus interval the fixation dot was blue instead of black to denote that this was the task-relevant interval. As described above, the stimuli in this interval either reversed in contrast polarity at 120Hz (invisible Kanizsa and control conditions) or the frames were interleaved with blank frames (visible Kanizsa, real luminance stimuli and control). During the stimulus presentation the small dark target dot also appeared at some location along the horizontal meridian. Its position was randomized to be either near the left or the right vertical boundary of the square region. The dot could either appear inside or outside the square region.

The dependent variable in this experiment was the distance between the target dot and the boundary of the square region and it was controlled by a 2-down, 1-up staircase procedure that converged on the threshold distance in each of the five experimental conditions at which performance was approximately 70.7% correct. That is, after every consecutive two correct trials the distance would decrease by one pixel (~0.03°) while after every incorrect trial it increased by one pixel. All five staircases started at a distance of 15 pixels (0.49°). The minimum and maximum that they could reach were one pixel and 25 pixels (0.82°), respectively.

Before the actual experiment we showed participants static examples of the Kanizsa and the control stimuli. Participants were instructed to judge whether the target dot was left or right of the (imaginary) vertical boundary of the square region. While showing them still images of the stimuli, we specifically explained to them that this boundary was defined by the exact center of the Gabor patches (i.e. the corner of the Pacman's mouth) and that this was identical in both the Kanizsa and the control conditions. We further informed them that there would be a third condition in which they should only see a subtly lighter grey square against the background but no Gabor patches. In order to become acquainted with the task, they then performed 1-2 blocks of the experiment with only the visible Kanizsa, the control and the real luminance condition. Finally, the actual experiment would commence. Participants were informed that the task was largely always the same as the familiarization run, although during this experiment there would be many more trials in which they either only see a light grey square but no Gabor patches or that they might even only see a grey blank screen. Except for the author, all participants were unaware of the experimental hypothesis. In debriefing none of the participants reported seeing any Gabor patches during the invisible condition.

Participants performed two consecutive runs comprising 500 trials each. These runs were further subdivided into 25 blocks. In every block each of the five conditions appeared 4 times in a pseudo-randomly interleaved order. Blocks were initiated by button-press. During block breaks a message on the screen reminded the participant of the task instructions and of the number of blocks they had already completed.

#### Data analysis

We determined the threshold distance in each of the five experimental conditions by calculating the mean distance across the final 15 reversals in the staircase procedure. Then we conducted a group level analysis in which we compared the average thresholds for conditions across participants. For this we used a two-way repeated-measures ANOVA with the factors visibility (visible vs invisible) and stimulus type (illusory vs control stimulus). As there was only one condition with a real luminance contour, the thresholds for this condition were not included in the ANOVA. However, we used paired t-tests to compare results for the illusory and control stimuli directly to the real luminance condition.

## Results

### Experiment 1

In this experiment, 17 participants judged the orientation (left vs right) of a triangle that was either defined by an illusory contour or a subtle real luminance edge. In a third control condition the Pacmen stimuli inducing illusory contours were rotated so that no triangle shape could be perceived.

#### Kanizsa Task

Accuracy across participants for discriminating the orientation of the triangle for the six conditions is displayed in Figure 4. We tested whether discrimination performance in each condition was above chance at the group level.

In the visible condition, the mean proportion of correct responses for the group was close to ceiling and clearly better than chance both for the real (*M*=0.97, *t*(16)=83.5, *p*< 0.001, *BF*_10_>9.8*10^18^) and the illusory condition (*M*=0.99, *t*(16)=120.8, *p*<0.0001, *BF*_10_>2.4*10^21^). In the control condition, however, the group performed significantly below chance level (*M*=0.30, *t*(16)=-3.87, *p*=0.001, *BF*_10_=29.3). There was considerable variability in performance for this condition ranging from 0 to 0.74.

In the invisible condition, only performance on the real triangle condition was significantly above chance level (*M*=0.71, *t*(16)=6.4, *p*<0.001, *BF*_10_=2608.7). Performance on both the illusory triangle condition (*M*=0.51, *t*(16)=1, *p*=0.332, *BF*_10_=0.385) and the control condition (*M*=0.50, *t*(16)=0.2, *p*=0.859, *BF*_10_=0.253) were at chance level. Importantly, performance on the real triangle condition was also significantly greater than for either the illusory contour (*t*(16)=6.1, p<0.0001, *BF*_10_=1475.4) or the control condition (*t*(16)=5.6, *p*<0.0001, *BF*_10_=688). In contrast, performance for the illusory contour did not differ from the control condition (*t*(16)=0.7, *p*=0.525, *BF*_10_=0.3).

#### Visibility Task

In a supplementary control task we asked participants to make a decision directly based on the inducers by reporting which one of the two bottom inducers contained the right angle. This assessed any residual perception of the inducer shapes under masking conditions. In the visible condition, the mean proportion of correct responses across participants was again close to ceiling and far above chance level (*M*=0.96, *t*(16)=22.2, *p*<0.001, *BF*_10_>2.9*10^10^). In contrast, participants' mean performance in the invisible condition did not significantly differ from chance level (*M*=0.48, *t*(16)=-1.3, *p*=0.227, *BF*_10_=0.489), suggesting that participants could generally not perceive the inducers under masking conditions. Note, however, that there was only anecdotal support for the null hypothesis (*BF*_10_>1/3).

The first experiment thus suggested that if participants were unaware of the inducers because they had been masked, they were unable to judge the orientation of the Kanizsa triangle. This supports the interpretation that illusory contours are not formed under these conditions.

**Figure 4.**
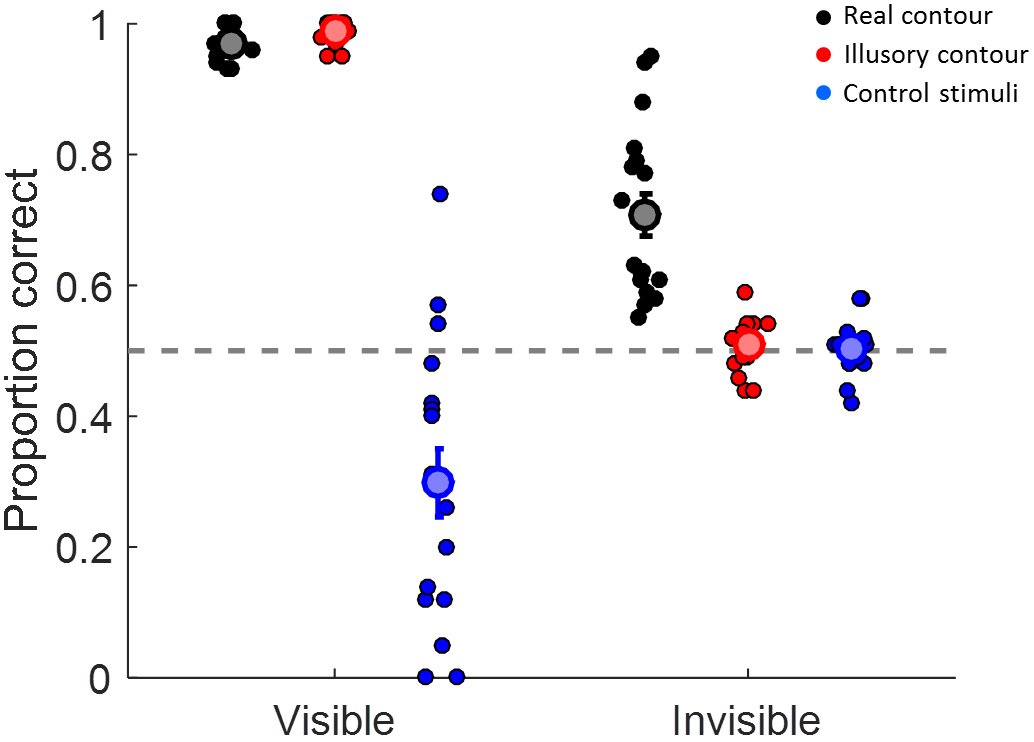
**Results of experiment 1**. Accuracy for discriminating the orientation of a triangle stimulus in visible or invisible (masked) trials. Each dot represents the performance of an individual participant in each of the conditions. The large symbols and error bars denote the mean ±1 standard error for each condition. Black: real luminance contour. Red: illusory (Kanizsa) contour. Blue: control stimuli.

### Experiment 2

In experiment 2 we changed the approach in a number of ways. First, instead of the brief stimulus presentations used in the first experiment we used counter-phase flicker to render stimuli invisible for prolonged periods. Moreover, we used an indirect measure of illusory contour processing: we took advantage of previous reports that the presence of an illusory contour stimulus enhanced participants' thresholds at discriminating the position of a small dot [23,39]. Here we tested whether this also occurred when the inducers generating the illusory contour were invisible. Because this task was more challenging than those in previous experiments, and to rule out that our previous results might have been due to insufficient practice or familiarity with psychophysical experiments, in this experiment we only tested a small group of well-trained psychophysics participants.

Figure 5 plots position discrimination thresholds in the different conditions for all individual participants and the group averages. Overall, thresholds measured while a Kanizsa stimulus was presented were significantly lower than those measured for control stimuli (*F*(1,4)=34.6, p=0.004). There was also a non-significant trend of lower thresholds during visible than invisible trials (*F*(1,4)=7.5, p=0.052). Importantly, there was a significant interaction between visibility and stimulus type (*F*(1,4)=15.9, p=0.016). In the visible condition thresholds measured while an illusory Kanizsa contour was present were significantly lower (*M*=0.08°, *t*(4)=-5.27, *p*=0.006, *BF*_10_=9.95) compared to the control condition without a contour (M=0.21°). In contrast, in the invisible condition the difference in thresholds for illusory contours (*M*=0.20°) was not significantly different from that for the control stimulus (*M*=0.21°, *t*(4)=-0.16, *p*=0.882, *BF_10_*= 0.4).

Thresholds measured for the real luminance contour were of a similar magnitude as those measured for the visible illusory contour (*M*=0.10°, *t*(4)=-0.92, *p*=0.409, *BF*_10_=0.55). In contrast, thresholds for the real luminance contour were significantly lower than for invisible Kanizsa stimuli (*t*(4)=3.84, *p*=0.019, *BF*_10_=4.53). Thresholds for the real luminance contour were also significantly lower than for either control stimulus (visible: *t*(4)=4.4, *p*=0.012, *BF*_10_=6.19; invisible: *t*(4)=5.3, *p*=0.006, *BF*_10_=9.9).

**Figure 5.**
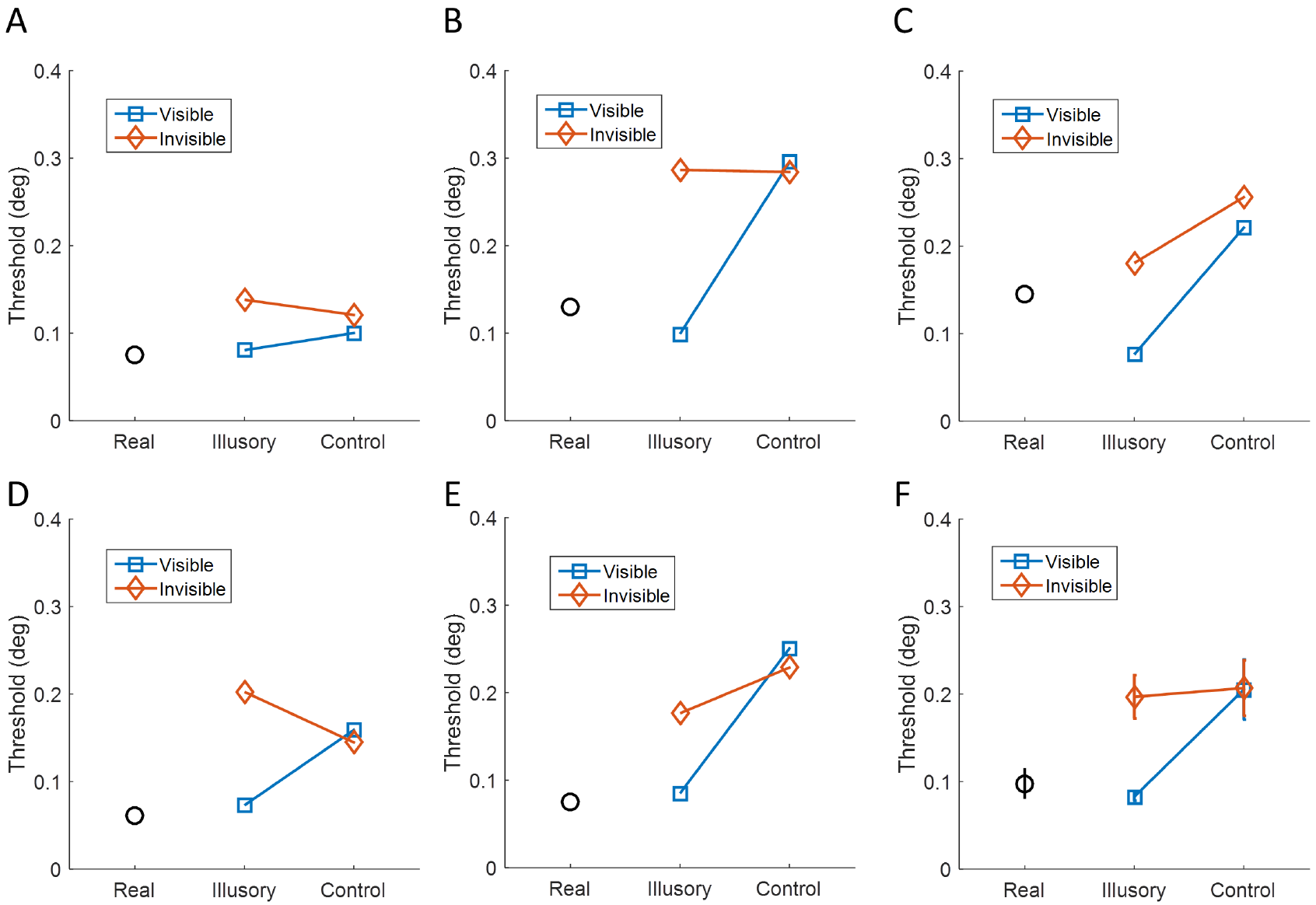
**Results of experiment 2**. Spatial discrimination thresholds per condition (real luminance vs illusory vs control) on a dot localization task while participants were presented with visible (blue squares), invisible (orange diamonds), or real luminance stimuli (black circles). A-E) Plots for the five individual participants. F) Thresholds averaged across participants ±1 standard error of the mean.

## Discussion

In two psychophysical experiments we tested whether the visual system forms illusory (Kanizsa) contours in the absence of awareness of the inducing context. Taken together, with previous experiments using dichoptic stimulation [22,23] all the findings suggest that illusory contours are not perceived when the inducers are masked.

In our first experiment, we directly measured illusory contour perception by asking participants to discriminate the orientation of a Kanizsa triangle. This procedure was akin to our earlier experiments [22] that showed no evidence either that illusory contours are formed when inducers are suppressed from awareness. In those experiments we employed CFS in which a dynamic, high-contrast mask presented to one eye suppresses awareness of a stimulus viewed by the fellow eye. Similarly, previous studies showed that illusory contours are not formed when the inducers are suppressed from awareness during binocular rivalry [23]. However, because previous reports indicate that illusory contours are processed by binocular neurons [30–33] the use of dichoptic stimuli may simply disrupt their processing.

Therefore here we used normal binocular viewing conditions and a different method to render the inducers invisible. In addition, we included a real luminance contour condition as a baseline check. This helped to rule out another trivial explanation for our earlier findings: The presence of the bright masks could possibly have obscured the detection of the subtle illusory boundary. Our real luminance contour was never masked because the masks only overlapped with the corners of the triangle; therefore, if participants were able to discriminate the subtle real luminance contour they should also be able to do so if an illusory contour were indeed formed. We only included participants for whom discrimination of this real luminance contour was significantly above chance levels. Nonetheless, participants were unable to discriminate the orientation of the illusory contour when inducers were masked. The phenomenological experience of the real luminance and the illusory contour is not perfectly identical (and it obviously is not for the visible conditions). However, the real luminance conditions were very faint while illusory contours tend to be subjectively quite salient. Thus it seems unlikely that participants should be able to detect only the real luminance contour but not the illusory contour.

Experiment 1 also contained control conditions in which the individual inducers had been rotated by 180° and thus no illusory triangle should be formed. These were essentially catch trials because participants should have been guessing, as there was no actual triangle to discriminate. This was clearly the case for control stimuli when inducers were invisible. However, for visible control stimuli discrimination performance varied widely from 0-0.74 proportion correct responses and was even significantly below chance level. The “correct” responses in this condition were dummy coded so that the stimulus-response mapping matched that for the equivalent illusory triangle stimuli before rotating the inducers. Therefore one inducer in the top row had a right-angled cutout that pointed inwards (Figure 1) and this would be consistent with the presence of a triangle oriented in the opposite direction (even though no actual triangle should be perceived). Some participants may have adopted this stimulus response mapping while others did not. It would therefore have been better to have all inducers point outwards in this control and randomize their locations. However, our actual results serendipitously support the fact that masking was effective in this experiment: The fact that participants performed at chance level when the control stimuli were masked rules out that they had any residual awareness of the inducers. The results of our explicit Visibility task after the main experiment also corroborated this conclusion.

The results of experiment 1 contradict previous claims that a “ghostly triangle” could be perceived when the inducers of a Kanizsa triangle were masked [24]. However, follow-up experiments failed to replicate these experiments but instead suggested that conscious processing of the inducers precedes awareness of the illusory contours and that these earlier results were in fact due to residual awareness of the inducers [45]. Another critical issue with these experiments is that the perception of illusory contours is based on a Yes or No judgment of whether a triangle shape was present. Such a task could theoretically be performed based on any residual awareness of the corners in the inducers. Our task required an explicit orientation discrimination of the illusory contour. If participants had been able to perform this task in spite of being unable to perceive the inducers, this would have been more conclusive evidence that illusory contours are indeed formed when inducers are masked.

Naturally, our design that split the main task from a test of awareness did not allow us to measure the awareness on each actual trial of the main experiment — only a dual-task design in which visibility is probed directly in the main experiment would permit this. However, a dual-task design cannot classify the awareness of individual trials perfectly as the participant's judgment of their own awareness is subject to variability. Dual-tasks also entail a division of attentional resources across the different task components. It could be argued that the Visibility task was more difficult than the Kanizsa task because the former required the judgment of one a small, peripheral feature of the stimulus. This may complicate the interpretation of chance performance on this task.

Experiment 1 also used very brief stimulus presentations and powerful, high-contrast masks to render stimuli invisible. Rather than the absence of awareness, the reason why invisible contours were not formed could be the brevity of the stimuli or because the mask fundamentally disrupted stimulus processing. This is certainly a possibility because previous research indicated that the processing of illusory contours occurs relatively slowly [36–38]. Therefore, in our second experiment, we used counter-phase flicker to mask the inducers instead of the masking methods employed in the earlier experiments. This allowed us to present stimuli for prolonged periods. While this is also a temporal masking procedure, the stimulus energy is constant during the entire presentation because only the polarity changes between frames. Furthermore, we exploited the fact that the presence of an illusory contour enhances performance on a dot localization task because it provides a visual aid for determining its spatial location [23,39]. We confirmed this advantage when inducers were not masked and performance was comparable to when we presented real luminance contours only. However, when inducers were rendered invisible this advantage for illusory contours disappeared. This task may actually be an even more appropriate test of the induction of illusory contours than testing discrimination of the contour itself as in experiment 1. For that experiment, the interpretation is unproblematic because we found no evidence of discrimination when inducers were masked. However, if we had found above chance performance for masked stimuli, it would have been impossible to conclude that this was not due to discrimination of the inducers. Pilot experiments for experiment 2 suggested that with the long-lasting counter-phase flicker masking, participants might in occasional trials have had some residual awareness of the corners in the inducers. Thus they might still have performed above chance at a shape/orientation discrimination on the purported illusory contour even though they did not in fact perceive any illusory contour. This problem also plagues many previous experiments that directly tested the presence of an illusory shape [36,37]. Therefore, only a task that exploits the presence of the illusory contour to modulate performance on an orthogonal task, like the dot localization task we used, can provide conclusive evidence that an illusory contour was in fact perceived. An alternative possibility could be a task that relies on a fine discrimination of a feature of the illusory contour such as its curvature [38] but even such discriminations may be confounded (with?) discrimination of the inducers [46,47].

In all tests of performance against chance levels we used Bayesian hypothesis tests that can quantify the strength of evidence for the null hypothesis indicating that participants were guessing [44]. None of these tests revealed *strong* evidence for the null hypothesis as typical Bayes Factors fell between 0.3-0.5. To establish more compelling support for the null hypothesis in those cases much larger samples would be required. Crucially, the Bayes Factor indicates by how much the observed evidence should update one's prior belief in the null or alternative hypothesis. Even if the evidence is relatively weak, any Bayes Factor below 1 is evidence *in favor of* the null and not for the alternative hypothesis. Unless one starts with a prior belief that people are clearly able to discriminate masked stimuli, even these modest Bayes Factors suggest that participants were probably guessing.

Several previous studies used stimulus manipulations that seek to disentangle the factors associated with Kanizsa-type illusory contours [39,48,49]. Rounding the corners of the Pacman inducers results in a notable reduction in the illusory contour percept and abolishes the concordant improvement on a spatial localization task. However, stimuli like this nonetheless activated higher extrastriate cortex to a similar degree as Kanizsa stimuli [39]. This is consistent with the theory that later stages of visual processing, presumably mediated by higher visual areas, segment surfaces and assign boundaries to objects. These segmentations are then fed back to early visual cortex to generate signals that are interpreted as illusory contours [48]. Practical support for this idea comes from transcranial magnetic stimulation experiments that disrupted neural processing either in object-sensitive lateral occipital (LO) cortex or in early visual areas V1 and V2 [50]. Critically, the disruption of LO cortex only abolished the illusory contour percept early after stimulus onset while disruption of early visual areas only did so at a later stage - presumably affecting feed-back signals rather than the early feed-forward response.

Additional stimulus processing that is unrelated to the actual formation of illusory contours could also explain previous reports that Kanizsa stimuli are faster to break through CFS masking than control stimuli [35]. The collinearity of edges and the thereby inferred surface may be processed even while the stimulus is suppressed from awareness — this may in turn produce an attentional signal that causes the stimulus to break suppression. Recent experiments that tested a range of visual control stimuli under CFS suggest that low-level properties of the stimulus determine the time it takes to break suppression [51]. Such stimulus-dependent effects are also plausible because attentional processing can occur without awareness of the stimulus [52–54]. However, this does not prove that any percept of an illusory contour was actually formed. This process may also explain why crowding interferes with discrimination of the inducer orientation but not with illusory contour formation [55].

While we did not manipulate the presence of illusory contours in this way in our experiments, we nonetheless controlled this factor by including real luminance contour conditions. In experiment 2 the presence of a contour should afford an improvement on an orthogonal spatial localization task. Such an improvement in localization thresholds only occurred for the real luminance contour or when Kanizsa inducers were visible. Improvements like this are not observed for stimuli that match the global characteristics but which do not produce illusory contours [39]. Therefore our results from this experiment strongly support the conclusion that illusory contours were simply not formed when inducers were masked.

Our results also agree with previous findings that only local processing occurs in the absence of awareness but that more complex analysis of scene geometry requires conscious processing [21]. This would also explain why the nature of stimulus representations in higher visual areas differs depending on awareness.

However, our findings do not accord with a number of other studies that suggest that illusory contours are processed unconsciously. Experiments on a patient with extinction due to a parietal lesion suggest that perception of a Kanizsa shape occurs even when some inducers are placed into parts of the visual field where the patient's conscious perception is impaired [26–29]. These findings indicate that visual processing operates at the surface-or object-based level. Segmenting and grouping the local features of an image into a coherent, global shape may only require awareness of some component features but it then spreads to the whole object. However, this is not sufficient evidence to demonstrate that these processes would occur when the participant is unaware of all components. More importantly, it also does not demonstrate that the percept of illusory contours was actually formed under these conditions but only that some processing of the features producing illusory contours under normal viewing conditions still occurred.

Another study used functional magnetic resonance imaging (fMRI) and an inattentional blindness paradigm to study illusory contours processing [49]. Participants were presented with a sequence of images, some of which contained Kanizsa shapes whilst others were various types of control stimuli. Simultaneously, participants were engaged in a demanding attentional task at fixation. A sub-group of participants subsequently reported not to have seen any Kanizsa stimuli. These participants nonetheless showed stronger fMRI responses to Kanizsa than control stimuli. Multivariate classification methods further demonstrated that the activation patterns produced by unseen Kanizsa stimuli were more reliable than those produced by control stimuli. The authors suggested that the neural signature of illusory contours differs from that of other, carefully matched stimuli and thus argued that illusory contours are processed even without awareness of the stimuli.

This finding is interesting because it tests the consequences of awareness without any experimental manipulation of the stimuli. The distinction of what is or is not processed without awareness only depends on the contents of the participant's consciousness. However, this also makes it difficult to interpret these results. First, it is unclear whether participants apparently oblivious to the presence of the stimuli really did not perceive any illusory shapes. Due to the design of the experiment, awareness could only be assessed after the main fMRI experiment rather than on a trial-by-trial basis (but see above our discussion why such trial-by-trial judgments of awareness are complicated). Only participants who reported having seen the actual Kanizsa stimulus during the experiment were classified as having had awareness of the stimuli. However, many of the candidate stimuli were similar Kanizsa shapes. Therefore it is possible that participants had some awareness of the stimuli, even if an imprecise one.

Second, while the control stimuli in the main fMRI experiment were very carefully matched to rule out the influence of global characteristics this can by definition only be an approximation: if conditions were perfectly matched, the stimulus would be identical and thus an illusory contour would be perceived. It is possible that the conditions resulting in an illusory contour percept are also particularly effective in producing discriminable activation patterns in visual cortex. For instance, the contrast energy along the mouths of the Pacmen inducers differed considerably between the Kanizsa and the global control stimuli. Thus, surface segmentation or collinear interactions may have also differed between these conditions.

All these issues again highlight an important point: The only way one can truly infer that illusory contours are formed is by using a measure that is specific to the presence of an illusory contour. This is indeed what we did in our previous CFS experiment [22], our present experiment 1, or what was done in several other previous experiments [23,34,50,55], by asking participants to directly report a feature of the contour. Another useful manipulation is the spatial localization task used by other studies [23,39] as well as our experiment 2. This is because the performance enhancement only occurs in the presence of an actual contour helping the participant to localize the target. Many previous studies suggesting that illusory contours are processed without awareness of the inducers are confronted with this problem. In the appendix we included another experiment in which we tested whether Kanizsa shapes masked from awareness could nevertheless provide an attentional cue for a subsequent visual search task. Even if a robust priming effect were found in such an experiment, this design simply cannot rule out alternative explanations. It only tests the consequences of the stimuli that may in fact not be specific to illusory contours.

In the same vein, neurophysiological and neuroimaging experiments have shown that illusory contours are encoded by neurons even in the early visual cortex [25,33,56–67]. The overwhelming majority of these experiments were conducted on awake participants. However, some neurophysiological studies reported neuronal tuning to illusory contours even in anesthetized animals [25,58,67]. Such findings seem to superficially contradict our conclusion that awareness of the inducing stimuli is necessary for the formation of illusory contours. However, these experiments are in fact a perfect illustration of the importance to distinguish between *perceptual experience* and correlated processing. Because animals are anesthetized in these experiments and thus by definition unaware of the stimuli, it is impossible to determine whether illusory contours were formed. The neural correlates of these stimuli could be related to the contextual processing of the inducing stimuli, such as the discontinuities detected by “end-stopped cells” or the detection of collinearity in the image. Such processing may indeed occur in the absence of awareness as is supported by our finding of collinear priming [21] and the induction of contextual visual illusions under masking conditions [2,3] or inattention [6]. Furthermore, it is quite likely that such stimulus processing is a necessary prerequisite for the formation of illusory contours. However, they do not conclusively prove that illusory contours are formed under anesthesia.

A related issue is that all of these studies use illusory contours induced by abutting lines (offset gratings) rather than Kanizsa shapes. While the two share a similar phenomenology, contours induced by abutting lines are arguably simpler and may be based mostly on local processing while Kanizsa shapes are likely to involve more complex inferences of surface depth and boundary ownership. Our experiments did not explicitly test illusory contours generated with abutting lines and therefore do not speak to the question whether such simpler illusory contours are in fact perceived when the inducers are masked.

We conclude that there is little evidence that Kanizsa-type illusory contours are processed when participants are not aware of the inducing context. This appears to be the case for a range of different methods to render the inducers invisible. However, all of these experiments employed physical stimulus manipulations, while it is likely that an experimental design that allows the use of the participant's own report to determine awareness on a trial-by-trial basis whilst also testing directly whether an illusory contour was in fact formed can answer this question conclusively.

## Acknowledgements

This research was supported by an ERC Starting Grant (310829) to DSS. We thank Pieter Moors, Jacob Jolij, and David Shanks as well as two anonymous reviewers for comments on a previous manuscript and Jonas Larsson for inspiring this study in the first place.

## Data and Materials

All stimulus code and data for this study are available for download at: https://dx.doi.org/10.6084/m9.figshare.2009952.

## Appendix Priming Experiment

Previous experiments used illusory contours to capture attention in a visual search task [68] or for priming [69,70]. In our priming experiment, we therefore tested if Kanizsa triangles rendered invisible using a similar masking procedure could be used as primes in a visual search task on a subsequent, visible stimulus array. <u>Like experiment 1</u>, this experiment comprised two consecutive tasks. The first, again called ‘Kanizsa’ task, was an indirect test that investigated whether priming with a Kanizsa triangle in the location of the target improves detection of the target. A second task consisted of a ‘Visibility’ task that directly tested the effectiveness of the masking technique.

## Study Design

In the context of this experiment, priming refers to the attentional cue provided by the prime shape about the location of the subsequently presented target in one of four possible locations that in turn affords a boost in behavioral performance on an orthogonal discrimination task. In the primed conditions the location of the target was always identical with the location of the prime, whereas in the unprimed conditions the prime shape was replaced by a control stimulus, which looked exactly like the distractors in the other three locations. Therefore, the unprimed condition provided a baseline that allowed the assessment of whether the triangle prime boosted the participants' performance by capturing their attention.

We tested priming with a 2×2×2 design with attentional priming (primed vs. unprimed), the type of priming triangle (real vs. illusory), and prime visibility (invisible vs. visible) as within subject factors. There were two different prime shapes: a real triangle defined by a strong luminance contrast or an illusory (Kanizsa) one. The visibility of the cue was manipulated by varying presentation times of the priming array that followed the mask. A short presentation time of 16.7ms was selected for the invisible conditions to render the prime invisible. In the visible conditions, however, the priming array was presented for 300ms, which left enough time for the prime to enter awareness.

Every participant completed two different tasks. Each task comprised 20 blocks of trials, where one block consisted of 32 trials. Across one task, the unprimed visible and invisible conditions appeared for a total of 160 times each, whereas the four primed conditions (real vs illusory and visible vs invisible) occurred 80 times each. Thus, half the block consisted of unprimed trials and the other half of primed ones.

Behavioral responses were given on a forced-choice discrimination task by button press (left or right arrow) on a standard computer keyboard. Each trial required either a left or a right response (see experimental procedure) and each response type appeared twice per block for each condition, except for the unprimed ones where it appeared four times per block. Conditions were selected pseudo-randomly for every trial but were counterbalanced within each block.

## Participants

Participants in this experiment were recruited among the UCL student population. Seventeen (13 female, age range: 21-32 years, mean age 24.7±2.5 years) normal, healthy participants took part in the experiment, including the two authors. All other participants were unaware of the experimental hypothesis.

## Stimuli

In half the trials, participants were cued with one of four possible cues. We created shape stimuli by placing four black discs (inducer elements, diameter=1.96°) in the configuration of a square (width=2.8°). In order to produce the illusory shape of a right isosceles triangle sitting on top of the square, a wedge was removed from each inducer element (Pacman). Wedges started at the center of their respective discs and extended to their edges. One wedge measured 90° and was always positioned either in the bottom left or bottom right inducer. The two wedges that formed the apexes of the triangle measured 45°. Finally, the wedge of the fourth inducer element measured 90°. This Pacman did not form part of the triangle and was positioned opposite the hypotenuse of the right isosceles triangle. It always faced the outside of the disc configuration. When the inducers were oriented with the gaps forming the three corners of a triangle, an illusory light grey Kanizsa triangle was visible on top of the black discs (Figure S1A, i and ii).

Real triangles were identical to the corresponding illusory shapes except that their contour was defined by a luminance contrast with a black line connecting the corners of the triangle (Figure S1A, iii and iv).

The control stimuli were created by altering the orientations of the three Pacmen by a systematic rotation of 180°. This modification broke the link between these inducers and thus no illusory triangle was perceived (Figure S1B). Control stimuli could also be horizontal mirror images of the illustrated examples.

Stimuli in the search array were right isosceles triangles with a contour defined by luminance (Figure S1C, i and ii). Target stimuli were triangles with the right angle pointing down, either to the left or to the right, while in distractors the right angle pointed up (Figure S1C, iii and iv). Participants were asked to detect the target and respond via button press whether its right angle was pointing to the left or to the right.

The mask consisted of a square of black lines connecting four discs, one at each corner. It was wide enough to cover all four Pacmen, as well as the location of the triangle within them (Figure S1D, i). In the ‘Visibility’ task, there was no search array. Instead, the stimulus array following the prime was a square containing a cross that gave no clue about the location and orientation of the target triangle (Figure S1D, ii).

The luminance of the background was 156 cd/m^2^. The luminance of inducer elements, the contour of the real triangle and the black of the mask was 2 cd/m^2^. Finally, the grey of the mask in the first task was 58.2 cd/m^2^.

Note that although there were differences between stimuli, the individual inducers remained the same and there was no change in the local image properties (i.e. square configuration of the four Pacmen and their position among the other 3 configurations). This enabled an assessment of the different effects of priming with the Kanizsa shape, as compared to the corresponding real triangle and ungrouped Pacmen or distractors in the unprimed conditions.

**Figure S1.**
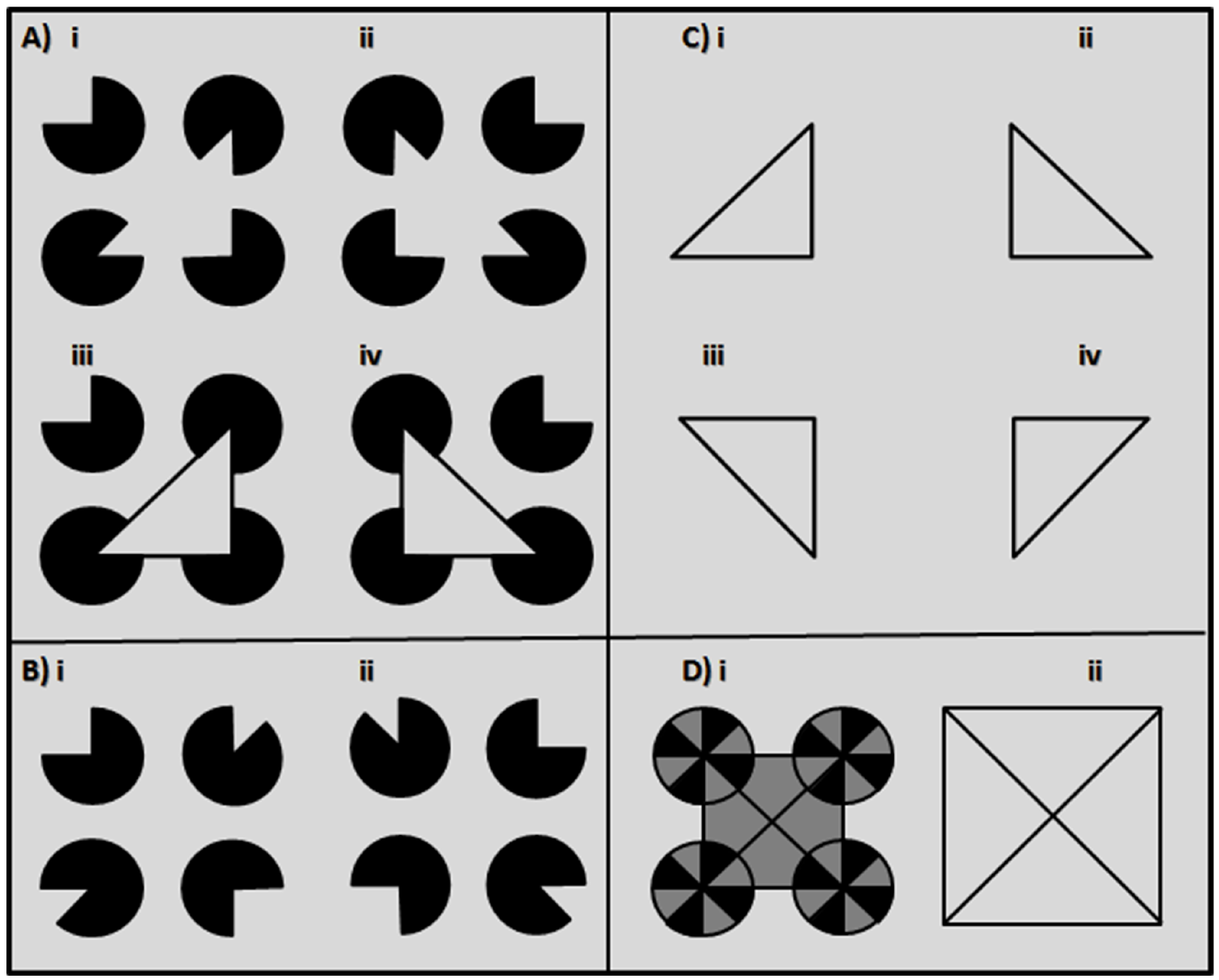
**Illustration of the stimuli in the priming experiment**. A) The four primes were formed of four inducer elements, of which three created either a real triangle or the illusory impression of a triangle. In half the trials, participants were cued with one of the four possible cues: (i) Kanizsa triangle pointing right, (ii) Kanizsa triangle pointing left, (iii) Real triangle pointing right, (iv)Real triangle pointing left. B) Examples of control stimuli used during cue presentation. Control stimuli were created by a systematic 180° rotation of the individual Pacmen, so that all gaps within the black discs pointed to the outside of the formation. Control stimuli could also be the horizontal mirror image of the presented examples. C) The two possible targets in the search task were right isosceles triangles with their right angle pointing down, either to the right (i) or to the left (ii). Distractor stimuli in the search array always pointed up, but varied between pointing left and right. D) (i) The mask used to render the prime invisible. (ii) The uninformative post-stimulus mask replacing the target array in the ‘Visibility’ task.

## Procedure

*Kanizsa task, indirect test assessing priming*:

At first, participants were instructed that they would see an array of four triangles of which only one, the target, had a right angle pointing down and were asked to decide whether this angle was pointing to the left or to the right by pressing the respective response keys.

The 20 experimental blocks were preceded by as many practice blocks as needed in order to fully familiarize the participants with the task requirements. Participants could initiate a new block by pressing any button on the keyboard and, if needed, take small breaks between blocks.

Participants were instructed to fixate a small black dot (0.2° wide) that was present in the center of the screen throughout the experiment. On each trial, the fixation dot was displayed alone for 500ms. Subsequently, a mask array was displayed for 100ms and was immediately followed by one of the priming arrays (see conditions in study design), which was presented for either 16.7ms or 300ms. This array could contain one of the two possible prime shapes (real or illusory triangle) or a simple control stimulus among the other three control stimuli. Immediately after this, a target array containing one target among three distractors was displayed and remained on the screen until the participant's response.

The response was self-paced; however, participants were instructed to respond as quickly as possible. Responses were made with the index (for left) and the middle or ring finger (for right), depending on how participants felt comfortable. For every response, the fixation dot provided feedback (100ms) by changing its color (green: correct; red: incorrect). Then, the next trial started without any delay. Figure S2A shows the general paradigm for the experimental procedure.

*Visibility task, direct test assessing prime discrimination:*

The second task assessed the effectiveness of the mask by measuring the conscious discriminability of the primes in a direct manner. For this purpose, participants were asked to make the same decision as in the previous task, but this time with respect to the prime. Thus, instead of discriminating whether the right angle of the target was pointing to the left or to the right, participants discriminated this aspect of the prime shape.

Procedure, stimuli and timing of the trial sequence in this task were kept identical to the previous indirect test, with a single exception: Instead of a target array, an array comprising new identical mask shapes covered the four stimuli locations (Figure S2B).

**Figure S2.**
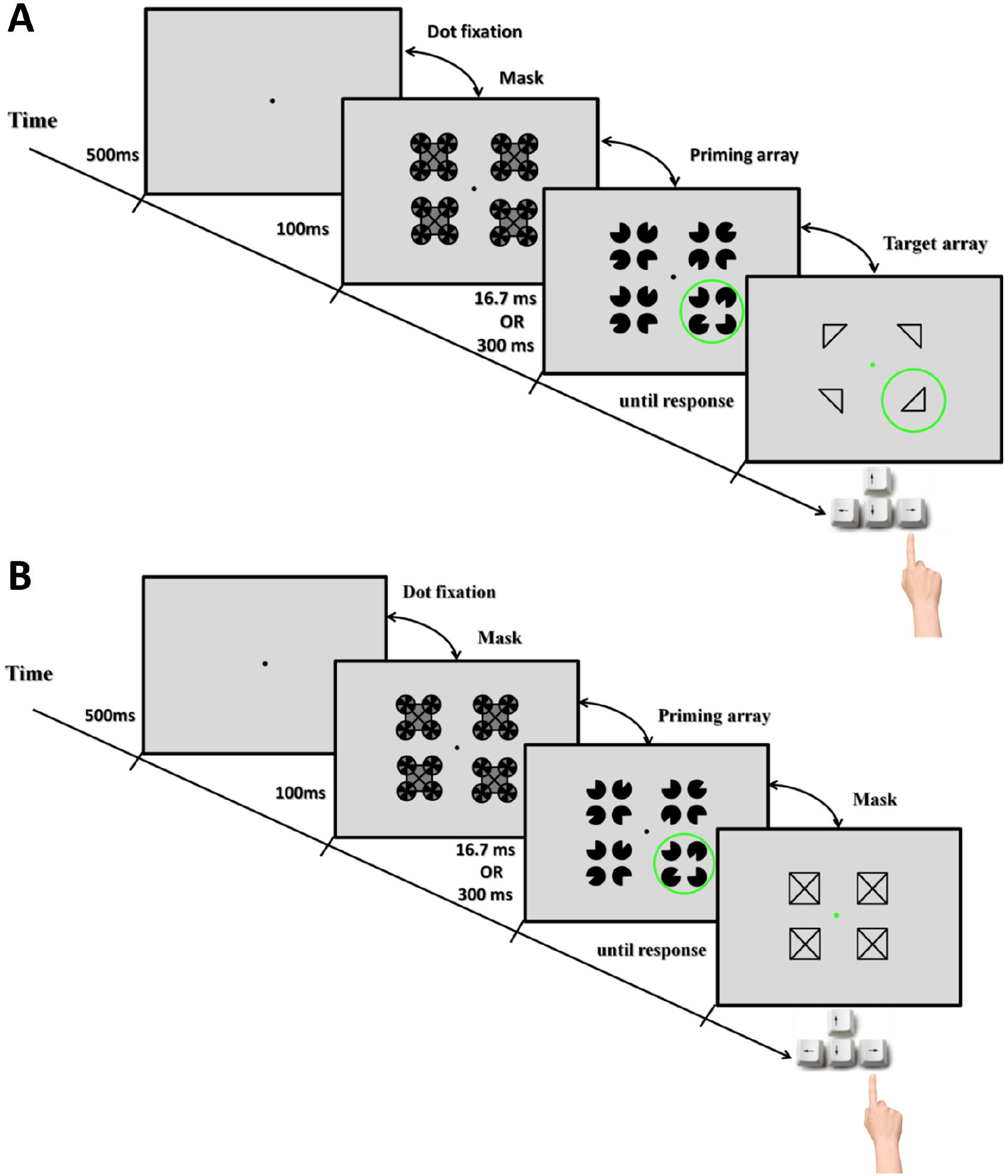
**Illustration of an example trial sequence in the priming experiment**. A) Kanizsa task: Each trial was composed of four frames: fixation period, mask, cueing array and target array. After the participant's response, the fixation point provided feedback for 100ms (green: correct; red: incorrect). The duration of each frame is shown on the time-line. The duration of the cueing array was varied between conditions: 16.7ms in the invisible conditions and 300ms in the visible ones. Note that in the unprimed conditions, the cueing array did not contain any prime shape and was composed exclusively of control stimuli. B) Visibility task: Trial procedure was identical to the one in the Kanizsa task except for the fourth frame in which the search array was replaced by an array of four identical mask shapes.

## Data Analysis

Response times (in milliseconds) and performance accuracy (in proportion correct) were measured for each of the six conditions. To analyze priming effects, response times were calculated on correct trials only.

For the Kanizsa task, accuracy of responses was less informative, as the target array was present until the participant's response. We reasoned that priming in the indirect Kanizsa task would be reflected by faster mean response times to primed than unprimed conditions. Therefore the priming effect was calculated by subtracting response times under priming from those without priming for correct responses.

Priming effects from all participants for all conditions were entered into a two-way repeated-measures analysis of variance (ANOVA) with prime visibility (visible vs invisible) and prime stimulus (real vs illusory triangle) as within subject factors. In addition to this, further comparisons between experimental conditions (visible or invisible/real or illusory) and control conditions (unprimed) were assessed with paired t-tests. We also determined the significance of the priming effect by comparing it against zero with one-sample t-tests.

In the visibility task, the variable of interest was the accuracy of responses as this allowed us to test whether participants could discriminate the orientation of the primes. One-sample t-tests were carried out at the group level for each condition individually in order to assess whether the participants' level of performance was significantly above chance (>0.5) and paired-tests were used to compare performance in different experimental conditions. As in experiment 1, for all t-tests we also calculated the default Bayes Factor in addition to frequentist statistics.

## Results

In the priming experiment we asked whether masked Kanizsa shapes could afford an attentional priming effect. We tested whether a Kanizsa stimulus presented as part of an array of four stimuli could prime participants to the location of a target in a visual search array presented subsequently. The prime could either be a Kanizsa triangle or a triangle defined by a real luminance contour. In the unprimed condition all four stimuli in the priming array were control stimuli without a triangle shape.

### Kanizsa Task

Overall participants made very few errors in determining whether the target pointed to the left or to the right. Indeed, the mean accuracy in all conditions was above 0.94. Therefore, for each experimental condition we quantified the effect of priming by the difference in response time with priming subtracted from that without priming on correctly answered trials (Figure S3A).

**Figure S3.**
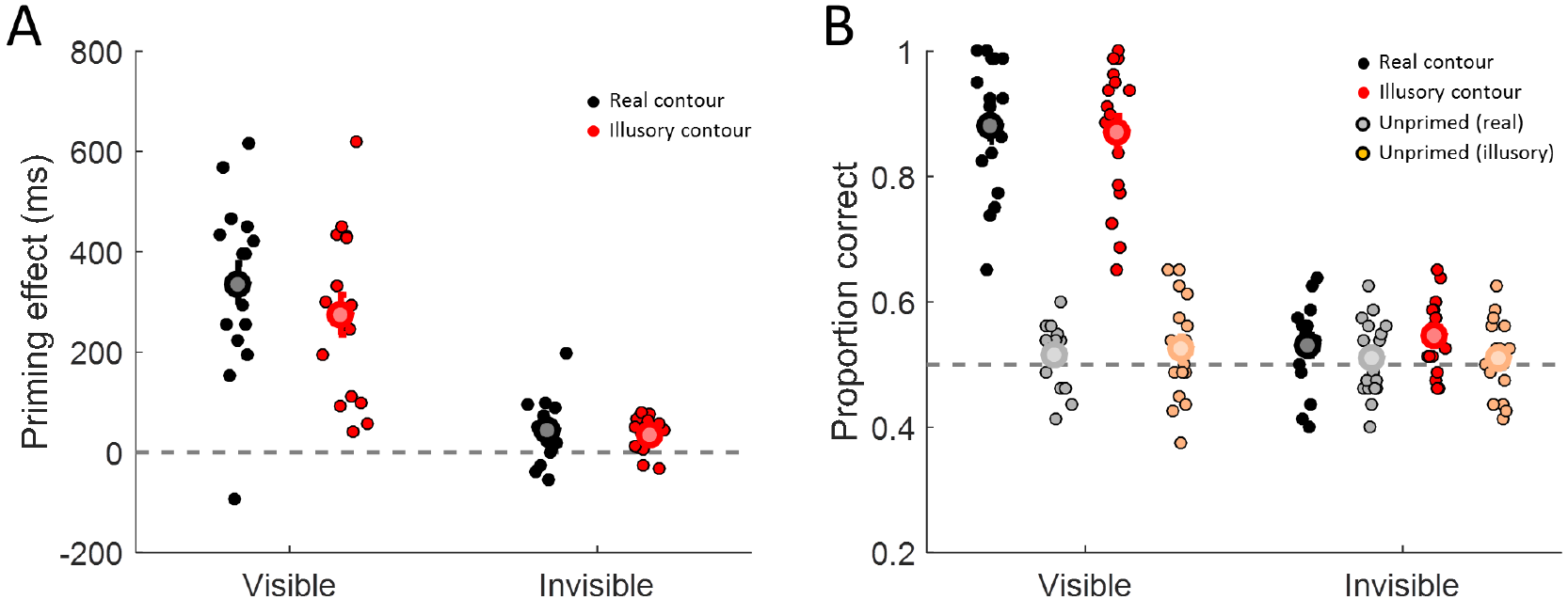
**Results of experiment 2**. Priming effect (response time for unprimed trials minus primed trials) on the Kanizsa task (A) and accuracy for discriminating the inducer location in the Visibility task (B) for visible or invisible (masked) trials. Each dot represents the mean performance of an individual participant in each of the conditions. The large symbols and error bars denote the mean ±1 standard error for each condition. Black: real luminance contour. Red: illusory (Kanizsa) contour. In B) Grey: unprimed trials dummy coded for real contour prime. Orange: unprimed trials dummy coded for illusory contour prime.

We conducted a two-way repeated-measures analysis of variance, with visibility (visible vs invisible) and stimulus (illusory vs real triangle) as within subject factors, to compare the priming effects (response time on unprimed minus primed trials) between conditions. There was a significant main effect of visibility of the prime (F(1,16)=60.06, p<0.001), and a significant main effect of stimulus type (*F*(1,16)=13.82, *p*=0.002). The interaction between visibility and the stimulus type was not significant (*F*(1,16=3.52, *p*=0.079).

There was a clear difference in the strength of priming between the visible and invisible condition. However, the critical test is whether the priming effect was significant, that is, if it differed from zero (i.e. a paired t-test between primed and unprimed response times). Unsurprisingly, the very pronounced priming effect for visible trials was extremely significant for both conditions (real: *M*=336ms, *t*(16)=8.2, *p*<0.001, *BF*_10_>4.1*10^4^; illusory: *M*=275ms, *t*(16)=6.9, *p*<0.001, *BF*_10_>5.8*10^3^). There was also a much more modest but nonetheless very significant priming effect for both stimulus types in the invisible condition when inducers were masked (real: *M*=43ms, *t*(16)=3, *p*=0.01, *BF*_10_=5.6; illusory: *M*=36ms, *t*(16)=4.4, *p*<0.001, *BF*_10_=80.8).

### Visibility Task

As in experiment 1 we used a direct Visibility test to assess whether participants might have had residual awareness of the priming stimuli under masking. The task procedure was the same as in the Kanizsa task except that the search array was replaced with a foil in which all four stimuli were identical and non-informative about the correct answer. We quantified the accuracy (proportion correct) with which participants could discriminate the orientation of the priming triangle (Figure S3B).

Unsurprisingly, in the visible condition performance for discriminating the orientation of a real (*M*=0.88, *t*(16)=14.9, *p*<0.001, *BF*_10_=9.5*10^7^) and an illusory triangle (*M*=0.87, *t*(16)=14, *p*<0.001, *BF*_10_=4.0*10^7^) was significantly above chance. Conversely, performance for the unprimed conditions, that is, stimulus arrays that contained no triangle shape, performance was consistently at chance (“real”: *M*=0.52, *t*(16)=1.3, *p*=0.21, *BF*_10_=0.52; “illusory”: *M*=0.53, *t*(16)=1.3, *p*=0.219, *BF*_10_=0.5; but note that these two conditions were dummy coded in this case as they contained the same stimuli).

Critically, performance in the invisible condition showed that discrimination in the primed with an illusory triangle condition was also significantly above chance (*M*= 0.55, *t*(16)=3.24, *p*=0.005, *BF*_10_=9.4), and discrimination of a real triangle showed a similar result even though it did not reach significance (*M*=0.53, *t*(16)=1.9, *p*=0.075, *BF*_10_=1.1). Performance for the two invisible triangles also did not differ significantly (*t*(16)=-0.9, *p*=0.385, *BF*_10_=0.35). Again, as expected both of the unprimed conditions were at chance level (“real”: *M*=0.51, *t*(16)=0.6, *p*=0.534, *BF*_10_=0.3; “illusory”: *M*=0.51, *t*(16)=0.7, *p*=0.494, *BF*_10_=0.31).

### Discussion

Taken together, the results of the priming experiment contradict those of experiment 1 and 2, as they suggest that Kanizsa triangles rendered invisible by masking could afford an attentional priming effect in a subsequent visual search task. However, a control experiment testing participants' ability to discriminate the prime shape orientation directly suggested that this small priming effect for invisible trials could have been due to some residual awareness of the masked triangle stimuli. When conducting such tests of awareness, it is of paramount importance to ensure that trials with masked and unmasked conditions are interspersed. When only masked conditions are tested for awareness, it is possible that the participant's performance is at chance even though there is in fact residual awareness in the main task due to what has been referred to as “priming of awareness” [71]. Conversely, the ability to correctly identify purportedly subliminal prime stimuli (as in our priming experiment) may have been enhanced by the inclusion of trials with clearly visible primes [72]. Either way, because we controlled for this possibility our visibility test should have provided a robust test of awareness. Therefore, the most parsimonious interpretation of the priming results is that participants had some residual awareness of the primes and that Kanizsa contours were probably not processed when inducers were masked. Using a different masking technique, for example the fast counter-phase flicker used in experiment 2 could be more effective and provide a better test of whether unconscious Kanizsa stimuli can act as attentional cues.

Critically, however, because priming experiments like this are only an indirect test of an effect that is not specific to illusory contour formation, this would not be informative. Even if we interpret the somewhat inconclusive results of this experiment as showing that masked triangles produced priming effects, this does not provide direct evidence that this advantage is actually caused by the presence of illusory contours. A Kanizsa triangle can clearly provide a salient attentional cue but this may be due to shared features, such as the collinearity of the edges in the inducers or the implication of a surface. This is also consistent with previous reports that priming of a shape discrimination task by subliminal stimuli depends on the strength of the salient region [73] and that salient regions of such stimuli are detected efficiently regardless of whether they are bounded by illusory contours [39,74]. Finding a priming effect thus only suggests that attentional cuing can occur without awareness but it does not rule out that other factors produced this attentional capture. Conversely, the absence of a priming effect would only confirm that whatever acts as attentional cue is disrupted. Therefore, the only meaningful test that illusory contours are indeed formed is via a measure that is *specific* to this percept.

